# E7-mediated repression of miR-203 promotes LASP1-dependent proliferation in HPV-positive cervical cancer

**DOI:** 10.1101/2024.01.08.574687

**Authors:** Molly R. Patterson, Aniek S. Meijers, Emma L. Ryder, James A. Scarth, Debra Evans, Amy L. Turner, Christopher W. Wasson, Janne E. Darell, Daisy Theobald, Joseph Cogan, Claire D. James, Miao Wang, John E. Ladbury, Iain M. Morgan, Adel Samson, Ethan L. Morgan, Andrew Macdonald

**Affiliations:** School of Molecular and Cellular Biology, Faculty of Biological Sciences, University of Leeds, Leeds, UK; Astbury Centre for Structural Molecular Biology, University of Leeds, Leeds, UK; Leeds Institute of Medical Research, St James’s University Hospital, University of Leeds, Leeds, UK; Leeds Institute of Rheumatic and Musculoskeletal Medicine, School of Medicine, University of Leeds, St-James University Teaching Hospital, Leeds, UK; Philips Institute for Oral Health Research, School of Dentistry, Virginia Commonwealth University (VCU), Richmond, Virginia, USA; VCU Massey Cancer Center, VCU, Richmond, Virginia, USA; School of Life Sciences, University of Sussex, Brighton, UK

## Abstract

Human papillomaviruses (HPV) are a major cause of malignancy, contributing to ∼5% of all human cancers worldwide, including most cervical cancer cases and a growing number of ano-genital and oral cancers. The major HPV viral oncogenes, E6 and E7, manipulate many host cellular pathways that promote cell proliferation and survival, predisposing infected cells to malignant transformation. Despite the availability of highly effective vaccines, there are still no specific anti-viral therapies targeting HPV or treatments for HPV-associated cancers. As such, a better understanding of viral-host interactions may allow the identification of novel therapeutic targets. Here, we demonstrate that the actin-binding protein LASP1 is upregulated in cervical cancer and significantly correlates with a poorer overall survival. In HPV positive cervical cancer, LASP1 depletion significantly inhibited proliferation *in vitro*, whilst having minimal effects in HPV negative cervical cancer cells. Furthermore, we show that the LASP1 SH3 domain is essential for LASP1-mediated proliferation in these cells. Mechanistically, we show that HPV E7 regulates LASP1 at the post-transcriptional level by repressing the expression of miR-203, which negatively regulated *LASP1* mRNA levels by binding to its 3’UTR. Finally, we demonstrated that LASP1 expression is required for the growth of HPV positive cervical cancer cells in an *in vivo* tumourigenicity model. Together, these data demonstrate that HPV induces LASP1 expression to promote proliferation and survival role in cervical cancer, thus identifying a potential therapeutic target in these cancers.

## Introduction

High-risk human papillomavirus (HR-HPV) infection is the underlying cause of almost all cervical cancers and several other anogenital and oropharyngeal cancers (1). These cancers are predominantly caused by HPV16 and HPV18, with 13 other HR-HPV types associated with cancer development (2). The drivers of HPV-induced proliferation are the viral oncogenes E5, E6 and E7 (3). During productive infection HPV E5 induces pro-proliferative EGFR signalling, promotes immune evasion and functions as a viral-encoded ion channel (4–7) However, HPV E6 and E7 are the primary oncogenes of viral transformation (3). The most well characterised function of these oncogenes is the inactivation of the p53 and pRb tumour suppressors, respectively (8–11); however, recent studies demonstrate that E6 and E7 modulate a plethora of host signalling pathways that have critical functions in cellular transformation (12–19).

The LIM and SH3 protein 1 (LASP1) was first identified in metastatic lymph nodes in breast cancer patients (20). The *LASP1* gene is located on chromosome 17q12 in humans and encodes a protein containing an N-terminal LIM domain followed by two actin-binding sites and a C-terminal SRC homology SH3 domain (21). This protein architecture allows LASP1 to engage in multiple protein–protein interactions, which may promote cell transformation. For example, the interaction of LASP1 with S100A11 promotes TGFβ-mediated epithelial-mesenchymal transition (EMT) in colorectal cancer (22). Furthermore, the binding of LASP1 to the tumour suppressor PTEN promotes PI3K/AKT signalling and tumour progression in nasopharyngeal carcinoma (23). Since its discovery, LASP1 has since been shown to be overexpressed in numerous cancers, including breast, lung, and colon cancer (24–29). Additionally, LASP1 expression is induced by several viruses that are associated with carcinogenesis, including Hepatitis C virus (HCV) and Hepatitis B virus (HBV) (30,31). However, whether LASP1 is modulated by HPV or in HPV-associated cancers is currently not known.

In this study, we demonstrated that LASP1 is upregulated in HPV positive (HPV+) cervical cancer and is significantly associated with worse overall survival. LASP1 is required for the proliferation of HPV+ cancer cells *in vitro* in an SH3-dependent manner, but is less important in HPV negative (HPV-) cancer cells. We further show that HR HPV induces LASP1 expression in both primary keratinocytes and cancer cell lines. Mechanistically, HPV E7 induces LASP1 expression by down-regulating the expression of miR-203, which directly targeted the *LASP1* 3’UTR. We further show that the tumour suppressive functions of miR-203 in cervical cancer cells is dependent on its targeting of LASP1. Finally, we demonstrated that LASP1 expression is required for the growth of HPV+ cervical cancer cells in an *in vivo* tumourigenicity model. Our findings suggest that LASP1 plays a key role in HPV-induced oncogenesis, highlight a potential therapeutic target in these cancers.

## Results

### LASP1 expression is increased in HPV positive cervical cancer

To investigate whether LASP1 played a role in HPV+ cervical cancer, we first utilised the GEO database. *LASP1* mRNA expression was upregulated in cervical cancer when compared with normal cervical tissue in several published datasets (Fig. 1A; (32–34)). An additional dataset demonstrated that *LASP1* mRNA expression was also higher in CIN3, a late, pre-malignant stage during the development of cervical cancer which represents severe dysplasia that may develop into cervical cancer (Fig. 1B; (35)). To confirm if LASP1 plays a role in the development of cervical cancer, we used cervical cytology samples from a cohort of HPV16+ patients and samples from healthy, HPV− patients as controls (19,36). Whilst *LASP1* mRNA expression was increased in all CIN stages, LASP1 protein was only significantly increased in CIN3 (Fig. 1C-D). These findings were corroborated by LASP1 immunofluorescence analysis in sections of cervical tissue from CIN1 and CIN3 samples (Fig. 1E). We next analysed LASP1 expression in a panel of cervical cancer cell lines, using primary normal human keratinocytes (NHKs) as a control. When compared with NHKs, both the mRNA and protein expression of LASP1 was significantly higher in HPV+ cervical cancer cells, with no significant difference observed between NHKs and HPV− C33A cervical cancer cells (Fig. 1F-G). To investigate this further, we utilised a tissue microarray (TMA) containing 9 normal cervical tissue samples and 38 cases of cervical cancer. LASP1 protein expression was significantly higher in cervical cancer when compared to normal cervical tissue, consistent with our cell line data (Fig. 1H). Finally, using the TCGA dataset, we observed that high LASP1 expression significantly correlated with worse overall survival in cervical cancer (Fig. 1I). Together, these data suggest that LASP1 may function as an oncogene in cervical cancer.

**Figure 1.**
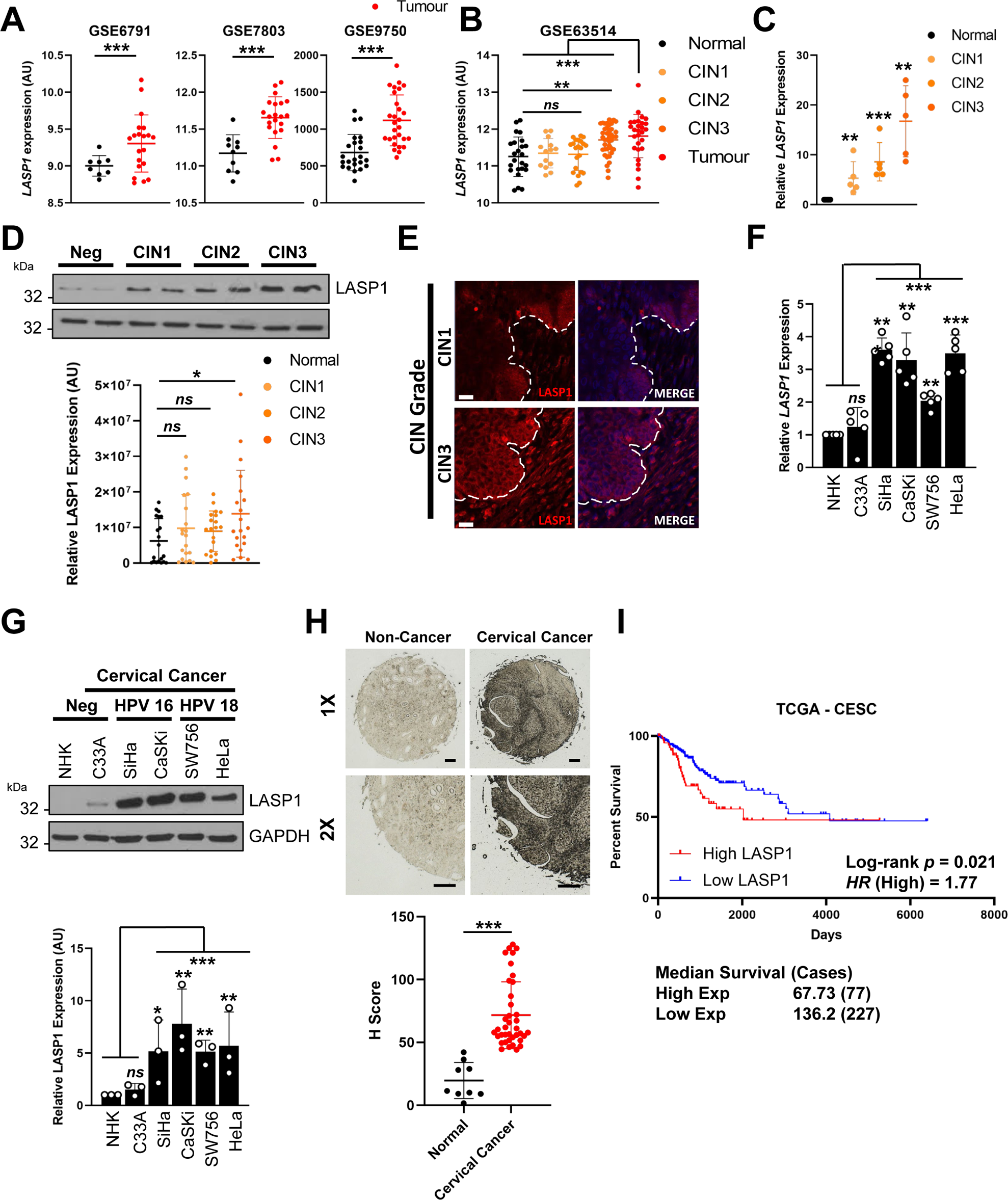
LASP1 expression is increased in HPV positive cancers. **A)** Scatter dot plot of *LASP1* mRNA expression in normal and cervical cancer tissue from the GEO database entries GSE6791, GSE7803 and GSE63514. **B)** Scatter dot plot of expression data acquired from the GSE63514 dataset. Arbitrary values for *LASP1* mRNA expression in normal cervix, CIN1 lesions, CIN2 lesions, CIN3 lesions and cervical cancer samples were plotted. **C)** RT-qPCR analysis of *LASP1* mRNA expression in a panel of cervical cytology samples representing normal cervical tissue (neg) and cervical disease of increasing severity (CIN 1 - 3) (*n* = 15 from each grade). Samples were normalised against *U6* mRNA expression. Data are displayed relative to neg samples. **D)** Representative western blot of LASP1 protein expression in a panel of cervical cytology samples representing normal cervical tissue (neg) and cervical disease of increasing severity (CIN 1 - 3). GAPDH was used as a loading control. Quantification from a larger samples set is shown below and data are displayed relative to NHK controls (*n* = 15 from each grade). **E)** Representative immunofluorescence analysis of tissue sections from cervical lesions of different CIN grades. Sections were stained for LASP1 expression (red) and nuclei were visualised with DAPI (blue). Images were acquired with identical exposure times and the white dotted line indicates the basal layer. Scale bar, 40 μm. **F)** RT-qPCR analysis of *LASP1* mRNA expression in HPV− normal human keratinocytes (NHK) and a panel of five cervical cancer cell lines – one HPV− (C33A), two HPV16+ (SiHa and CaSKi), and two HPV18+ (SW756 and HeLa). Samples were normalised against *U6* mRNA expression. Data are displayed relative to NHK controls. **G)** Representative western blot of LASP1 protein expression in HPV− normal human keratinocytes (NHK) and a panel of five cervical cancer cell lines. GAPDH was used as a loading control. Quantification from a larger samples set is shown below and data are displayed relative to NHK controls. **H)** Representative immunohistochemistry analysis and scatter dot plots of quantification of normal cervical (*n* = 9) and cervical cancer (*n* = 38) tissue sections stained for LASP1 protein. Scale bar, 50 μm. **I)** Overall survival analysis of cervical cancer data based on *LASP1* mRNA expression. Survival data were plotted using the Kaplan–Meier survival curve from 297 cases. Red indicates high expression, cyan indicates low expression. Error bars represent the mean +/− standard deviation of a minimum of three biological repeats unless otherwise stated. *ns –* not significant, \**p* < 0.05, \*\**p* < 0.01, \*\*\**p* < 0.001 (Student’s *t*-test).

### HPV E7 increases LASP1 expression

As our data demonstrated that LASP1 was increased in HPV+ cervical cancer cell lines, but not in an HPV− cervical cancer cell line, we investigated if HPV played an active role in the upregulation of LASP1. Firstly, we analysed the expression of LASP1 in NHKs stably harbouring the HPV18 genome (6,37). When compared to NHKs, *LASP1* mRNA expression was ∼3 fold higher in HPV18 containing keratinocytes (Fig. 2A). In line with this, LASP1 protein expression was also higher in NHKs containing the HPV18 genome from two individual donors (Fig. 2B). This was also confirmed for another HR-HPV type, HPV16, from two different clonal cell lines (Fig. 2C; (36)).

**Figure 2.**
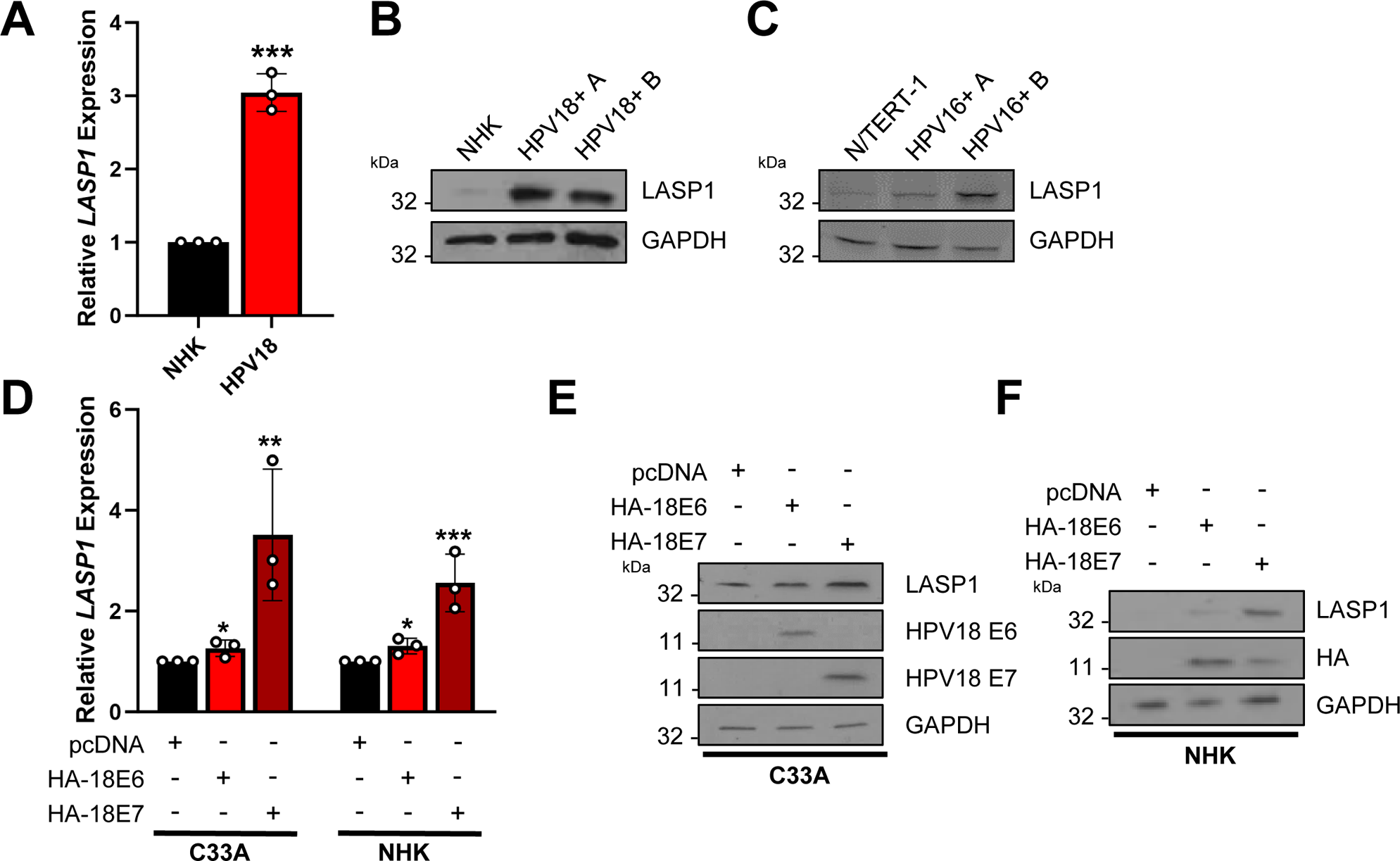
HPV E7 increases LASP1 expression. **A)** RT-qPCR analysis of *LASP1* mRNA expression in NHKs and HPV18 containing NHKs. Samples were normalised against *U6* mRNA expression. Data are displayed relative to NHK controls. **B)** Representative western blot of LASP1 protein expression in NHKs and HPV18 containing NHKs. Data from two individual HPV18 containing NHK clones (A and B) is shown. GAPDH was used as a loading control. **C)** Representative western blot of LASP1 protein expression in N/TERT-1 and HPV16 containing N/TERT-1. Data from two individual HPV16 containing N/TERT-1 clones (A and B) is shown. GAPDH was used as a loading control. **D)** RT-qPCR analysis of *LASP1* mRNA expression in HPV− C33A cells after transfection with HPV18 E6 or E7 for 48 hours. Samples were normalised against *U6* mRNA expression. Data are displayed relative to cells transfected with pcDNA3.1 as a vector control. **E**-**F)** Representative western blot of LASP1 protein expression in HPV− C33A cells **(E)** and NHKs **(F)** after transfection with HPV18 E6 or E7 for 48 hours. Lysates were probed for the expression of LASP1, HPV18 E6 and HPV18 E7 (C33A) or LASP1 and HA (NHK). GAPDH was used as a loading control. Error bars represent the mean +/− standard deviation of a minimum of three biological repeats unless otherwise stated. *ns –* not significant, \**p* < 0.05, \*\**p* < 0.01, \*\*\**p* < 0.001 (Student’s *t*-test).

The HR-HPV oncogenes E6 and E7 are the primary drivers of tumourigenesis in HPV− associated cancers. We therefore investigated if E6 or E7 play a role in the increased LASP1 expression in HPV containing cells. To investigate this, we over-expressed HPV18 E6 and E7 in HPV− C33A cells and in NHKs. Although HPV E6 resulted in a slight increase in *LASP1* expression at the mRNA level, HPV E7 resulted in a significant increase at both the mRNA and protein level in both cell lines (Fig. 2D-F). These results demonstrate HPV E7 is primarily responsible for the high expression of LASP1 observed in HPV+ cervical cancer and HPV containing keratinocytes.

### LASP1 expression is regulated by miR-203 in an E7-dependent manner

HPV E6 and E7 manipulate microRNA (miRNA) networks in order to control host gene expression (16,38–40). Furthermore, LASP1 has previously been shown to be regulated by several miRNAs in diverse cancers (25,41–45). Of these, we and others have previously shown that miR-203 is downregulated in cervical cancer (40,46–48). To understand if miR-203 regulates LASP1 in cervical cancer, we first analysed the panel of cervical cytology samples and observed a significant inverse correlation between miR-203 levels and *LASP1* mRNA expression (Fig. 3A). To identify if there was a functional relationship between miR-203 and LASP1, cells were transfected with an miR-203 mimic and *LASP1* mRNA levels were assessed by RT-qPCR. miR-203 expression led to a dose-dependent decrease in *LASP1* mRNA and protein levels in HeLa and SiHa cells (Fig. 3B-C). To confirm if *LASP1* is a direct target of miR-203, we identified the miR-203 binding site in the *LASP1* mRNA 3’UTR (using the AceView program) and generated a mutant 3’UTR sequence lacking complementarity to miR-203 (Fig. 3D). HeLa and SiHa cells were transfected with the miR-203 mimic and a reporter plasmid containing a luciferase sequence fused to either the wild type (WT) or mutant *LASP1* 3’UTR. Cells transfected with the miR-203 mimic showed decreased luciferase levels, indicating that miR-203 directly targets the *LASP1* 3’ UTR (Fig. 3E). In contrast to the WT, luciferase expression from the mutant *LASP1* 3’ UTR plasmid was unaffected by the miR-203 mimic (Fig. 3E).

**Figure 3.**
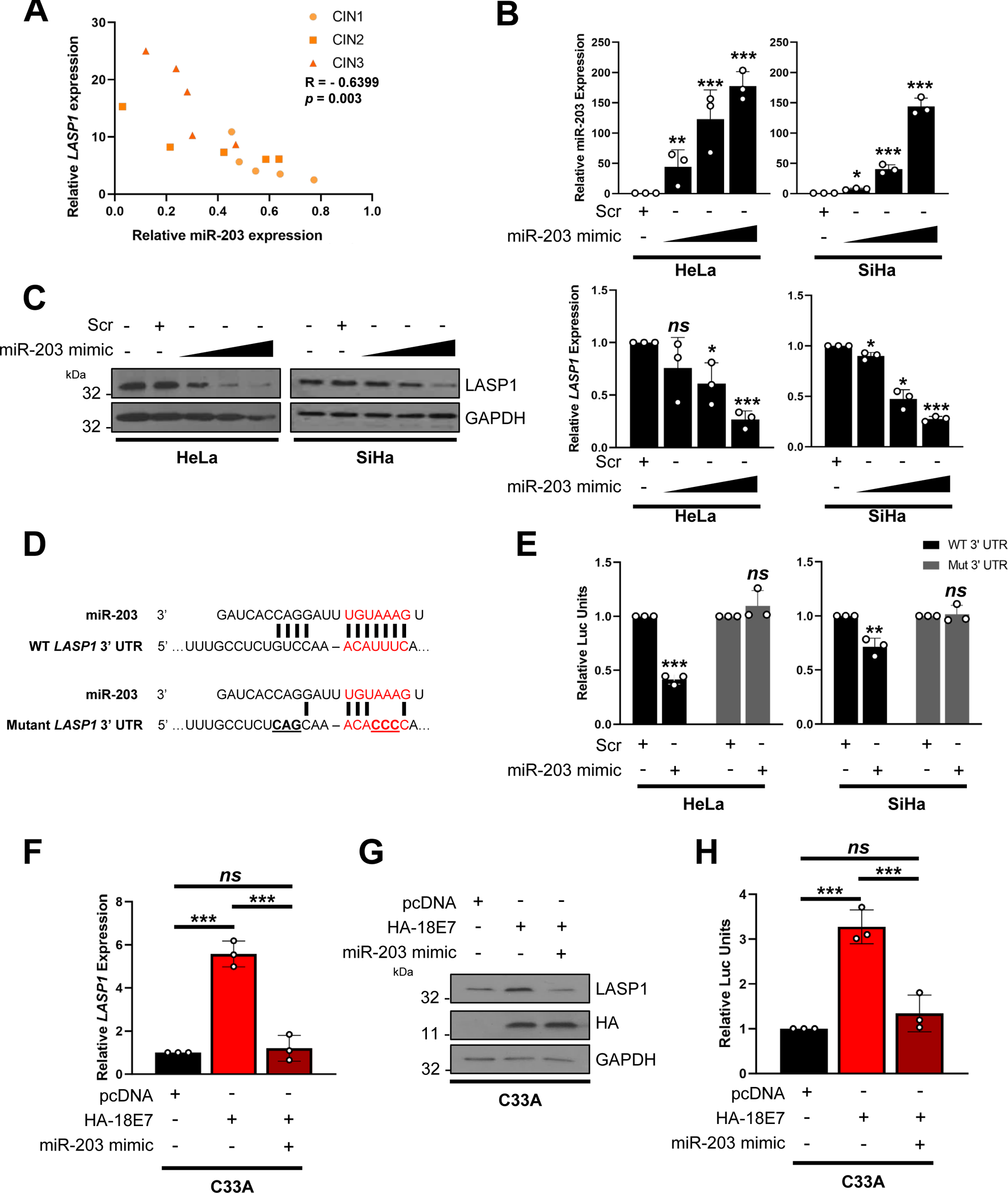
LASP1 expression is regulated by miR-203 in HPV+ cervical cancer. **A)** A) Graph showing correlation between miR-203 and *LASP1* mRNA expression in matched patient samples of cervical disease (n = 5 from each grade). **B)** (top) miScript analysis of miR-203 expression in HeLa and SiHa cells after transfection of increasing concentration of a miR-203 mimic. (bottom) RT-qPCR analysis of *LASP1* mRNA expression in HeLa and SiHa cells after transfection of increasing concentration of a miR-203 mimic. Samples were normalised against *snORD68* expression (miR-203) or *U6* mRNA expression (*LASP1*). Data are displayed relative to a scramble control. **C)** Representative western blot of LASP1 protein expression in HeLa and SiHa cells after transfection of increasing concentration of a miR-203 mimic. GAPDH was used as a loading control. **D)** Schematic of *LASP1* 3’UTR showing miR-203 binding site and mutant miR-203 binding site. **E)** Luciferase reporter assays from HeLa and SiHa cells co-transfected with miR-203 mimic and either a wild-type *LASP1* 3’UTR reporter plasmid or a mutant plasmid that lacks the putative miR-203 binding site. Data presented are relative to an internal firefly luciferase control. **F)** RT-qPCR analysis of *LASP1* mRNA expression in C33A cells after transfection with HPV18 E7, with or without a miR-203 mimic. Samples were normalised against *U6* mRNA expression. Data are displayed relative to cells transfected with pcDNA3.1 as a vector control. **G)** Representative western blot of LASP1 protein expression in C33A cells after transfection with HPV18 E7, with or without a miR-203 mimic. GAPDH was used as a loading control. **H)** Luciferase reporter assays from C33A cells co-transfected with HPV18 E7 and a wild-type *LASP1* 3’UTR reporter plasmid, with or without a miR-203 mimic. Data presented are relative to an internal firefly luciferase control. Error bars represent the mean +/− standard deviation of a minimum of three biological repeats unless otherwise stated. *ns –* not significant, \**p* < 0.05, \*\**p* < 0.01, \*\*\**p* < 0.001 (Student’s *t*-test).

Next, we wanted to confirm if HPV E7 mediated LASP1 upregulation requires the repression of miR-203 expression. In order to do this, we expressed HPV18 E7 in C33A cells. HPV E7 expression increased *LASP1* mRNA expression, LASP1 protein expression and luciferase levels driven by the WT *LASP1* 3’ UTR luciferase plasmid (Fig. 3F-H); however, co-transfection of E7 and a miR-203 mimic ablated this increase. Taken together, these data suggest miR-203 directly targets *LASP1* through binding to the miR-203 binding site within the *LASP1* 3’ UTR and that HPV E7-mediated LASP1 expression is miR-203-dependent.

### LASP1 promotes proliferation in HPV+ cervical cancer cells in an SH3-dependent manner

To investigate the role of LASP1 in cervical cancer, we first depleted LASP1 using a pool of four specific siRNA. Transfection of these siRNAs resulted in a significant loss of LASP1 protein expression in HPV+ HeLa and SiHa cells and HPV− C33A cells (Supp. Fig. 1A and 2A). LASP1 depletion led to a similar decrease in HPV E6 and E7 expression. Next, we investigated the impact of LASP1 depletion on cervical cancer proliferation. In both HPV+ cell lines, LASP1 depletion resulted in a significant reduction cell growth (Supp. Fig. 1B) and colony formation under anchorage dependent (Supp. Fig. 1C) and independent conditions (Supp. Fig. 1D). In contrast, depletion of LASP1 had minimal impact on cell proliferation or colony formation in HPV− cervical cancer cells (Supp. Fig. 2B-D).

LASP1 is comprised of two well characterised protein domains, the LIM domain and the SH3 domain (Supp Fig. 3A; (49)). We investigated the role of each domain in the proliferation phenotypes observed in cervical cancer cells. Expression of WT LASP1 enhanced cell growth and colony formation in HPV− C33A cells (Supp. Fig. 3B-E). We then expressed LASP1 mutants in which either the LIM domain or SH3 domain were deleted (GFP-LASP1ΔLIM or GFP-LASP1ΔSH3). Deletion of the LIM domain enhanced cell growth and colony formation at a similar level to WT LASP1; however, deletion of the SH3 domain failed to enhance cell growth or colony formation, suggesting that this domain was essential for the increased proliferation phenotypes observed upon LASP1 expression. To confirm these data, we generated two individual monoclonal HeLa cell LASP1 knock down cells using shRNAs (LASP1 knockdown (KD)1 and LASP1 KD4). As observed with transient LASP1 depletion, LASP1 KD reduced HPV E6 and E7 expression and significantly impaired cell growth and colony formation (Fig. 4A-D). We then complemented our LASP1 KD cells with our LASP1 mutants (Fig. 4E). WT LASP1 and the LIM domain mutant fully restored cell growth and colony formation to control levels (Fig. 4F-H). However, the SH3 domain mutant failed to restore the proliferation defects observed upon LASP1 KD. These data demonstrate that LASP1 promotes proliferation in HPV+ cervical cancer cells and this is dependent on the SH3 domain of LASP1.

**Figure 4.**
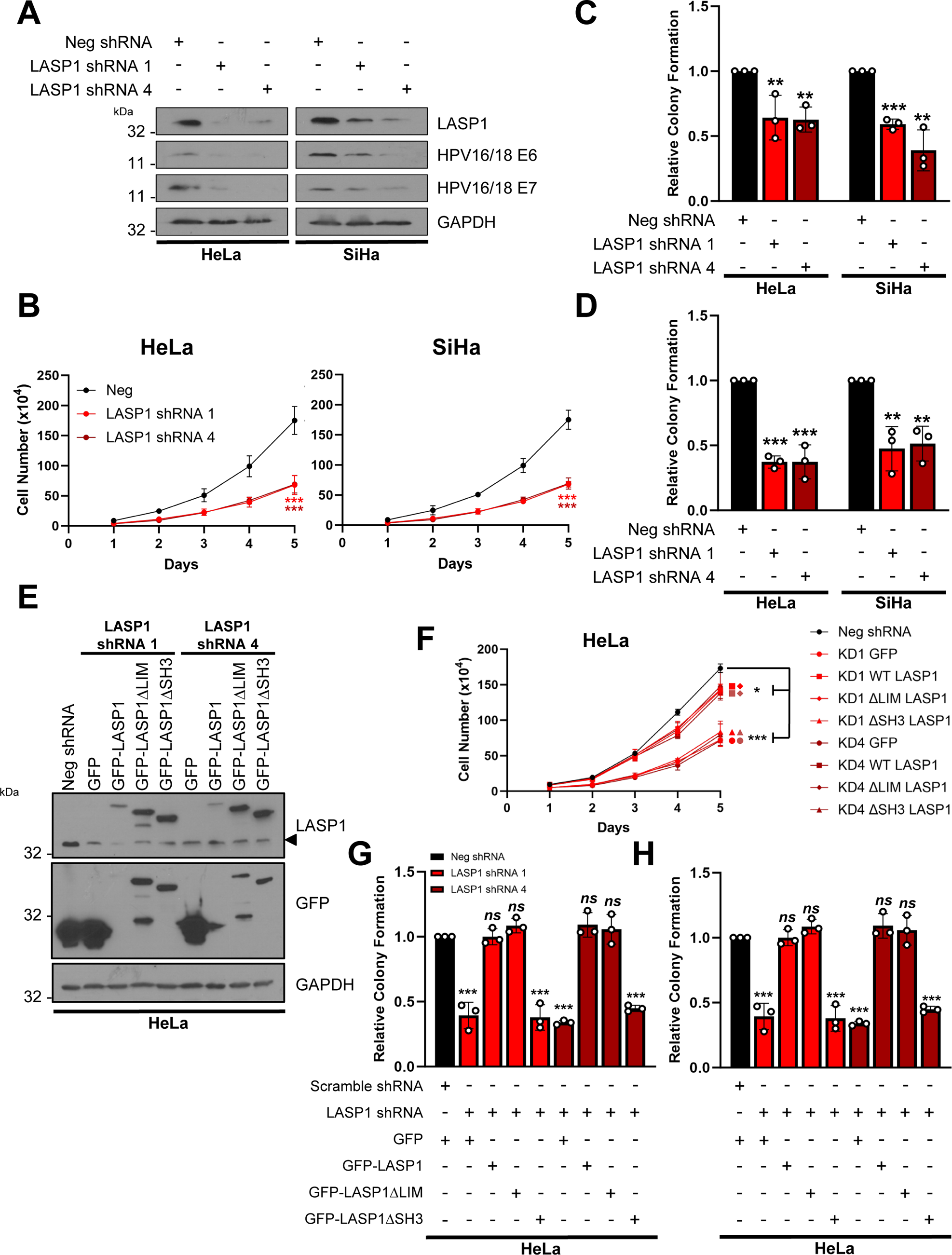
LASP1 promotes proliferation in HPV+ cervical cancer cells in an SH3-domain dependent manner. **A)** Representative western blot of LASP1 protein expression in HeLa and SiHa cells after shRNA mediated depletion of LASP1 (LASP1 knock down (KD) cells). Two monoclonal populations are shown for each cell line. Lysates were probed for LASP1, HPV E6 and HPV E7. GAPDH was used as a loading control. **B)** Growth curve assay in HeLa and SiHa LASP1 KD cells. **C)** Colony formation assay in HeLa and SiHa LASP1 KD cells. **D)** Soft Agar assay in HeLa and SiHa LASP1 KD cells.**E)** Representative western blot of LASP1 mutants in HeLa and SiHa LASP1 KD cells. GFP-LASP1, GFP-LASP1ΔLIM and GFP-LASP1ΔSH3 expression were confirmed using GFP and LASP1 antibodies. Arrow indicates endogenous LASP1 expression. GAPDH was used as a loading control. **F)** Growth curve assay in HeLa LASP1 KD cells and HeLa LASP1 KD cells expressing GFP-LASP1, GFP-LASP1ΔLIM and GFP-LASP1ΔSH3. **G)** Colony formation assay in HeLa LASP1 KD cells and HeLa LASP1 KD cells expressing GFP-LASP1, GFP-LASP1ΔLIM and GFP-LASP1ΔSH3. **H)** Soft Agar assay in HeLa LASP1 KD cells and HeLa LASP1 KD cells expressing GFP-LASP1, GFP-LASP1ΔLIM and GFP-LASP1ΔSH3. Error bars represent the mean +/− standard deviation of a minimum of three biological repeats unless otherwise stated. *ns –* not significant, \**p* < 0.05, \*\**p* < 0.01, \*\*\**p* < 0.001 (Student’s *t*-test).

### miR-203 inhibits the proliferation of HPV+ cervical cancer cells via LASP1 repression

To determine if the tumour suppressive effect of miR-203 is due to the repression of LASP1, we restored LASP1 expression in HeLa and SiHa cells expressing a miR-203 mimic (Fig. 5A and B). As we previously demonstrated, miR-203 significantly reduced cell growth and colony formation in HPV+ cervical cancer cells (Fig. 5C-E; (40)). Restoration of LASP1 in miR-203 mimic expressing cells completely abolished the proliferation defects observed, demonstrating that LASP1 is a major target of miR-203 in HPV+ cervical cancer cells.

**Figure 5.**
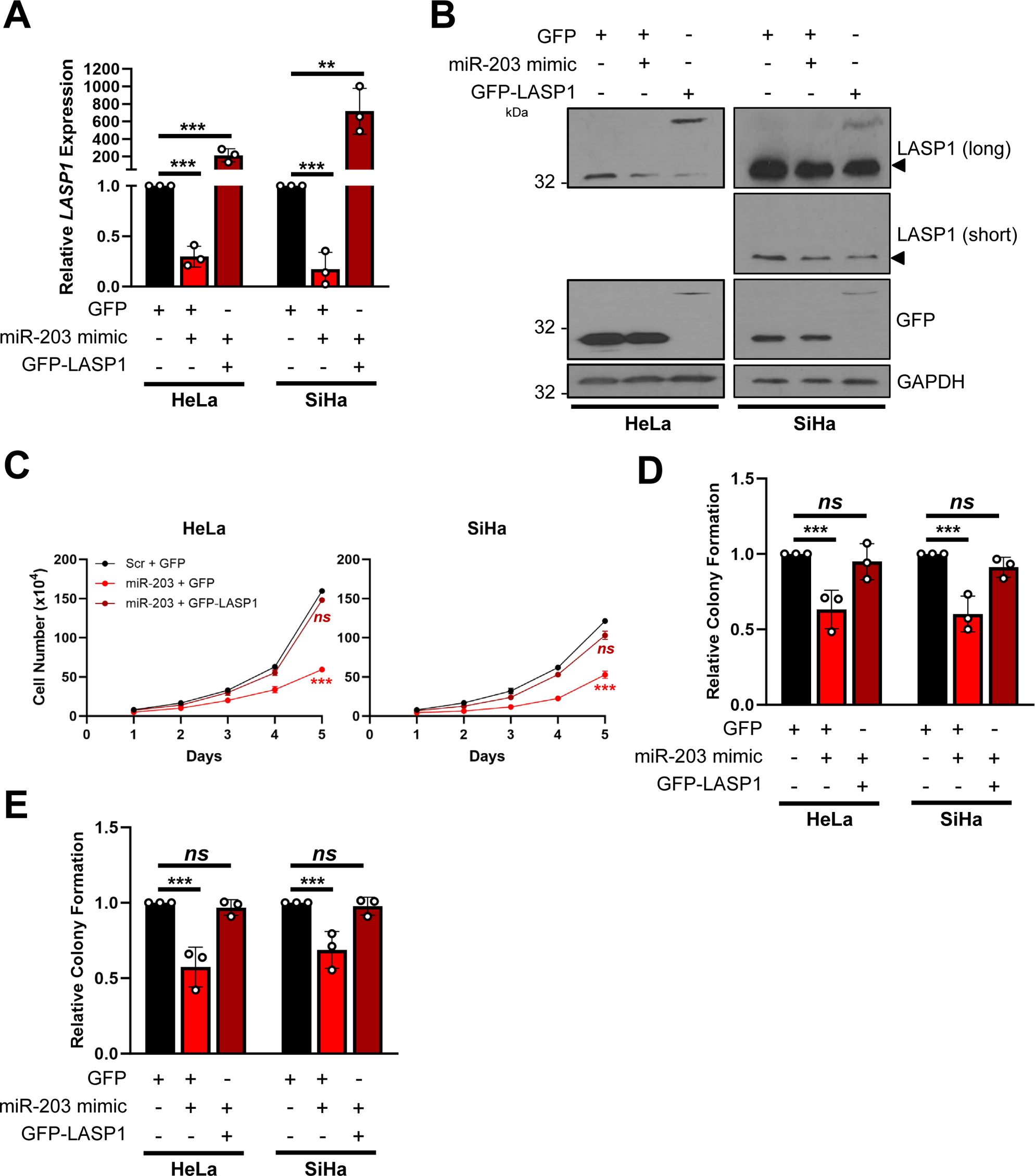
miR-203 inhibits the proliferation of HPV+ cervical cancer cells via LASP1 repression. **A)** RT-qPCR analysis of *LASP1* mRNA expression in HeLa and SiHa cells after transfection of miR-203 mimic, with or without GFP-LASP1. Samples were normalised against *U6* mRNA expression. Data are displayed relative to a GFP control. Representative western blot of LASP1 protein expression in HeLa and SiHa cells after transfection of miR-203 mimic, with or without GFP-LASP1. Lysates were probed for LASP1 and GFP. GAPDH was used as a loading control. **C)** Growth curve assay in HeLa and SiHa cells after transfection of miR-203 mimic, with or without GFP-LASP1. **D)** Colony formation assay in HeLa and SiHa cells after transfection of miR-203 mimic, with or without GFP-LASP1. **E)** Soft Agar assay in HeLa and SiHa cells after transfection of miR-203 mimic, with or without GFP-LASP1. Error bars represent the mean +/− standard deviation of a minimum of three biological repeats unless otherwise stated. *ns –* not significant, \**p* < 0.05, \*\**p* < 0.01, \*\*\**p* < 0.001 (Student’s *t*-test).

### LASP1 depletion impairs HPV+ cervical cancer cell growth in an *in vivo* tumourigenicity model

To confirm our *in vitro* observations, we performed *in vivo* tumourigenicity experiments using SCID mice. Animals were subcutaneously injected with HeLa Neg shRNA or HeLa LASP1 KD4 cells. Tumour development was monitored, revealing rapid growth in the Neg shRNA control group (Fig. 6A). However, in LASP1 KD tumours, growth was significantly delayed. To assess this quantitively, the period of time between injection of tumours and growth to a set volume (400 mm^3^) was calculated. This demonstrated that LASP1 KD tumours took an additional 18 days on average to reach an equivalent size (Fig. 6B). Further, animals bearing LASP1 KD tumours displayed significantly prolonged survival (Fig. 6C; Neg shRNA – median survival of 37 days, LASP1 KD – median survival of 72 days). Together, these data demonstrate that LASP1 is a critical driver of HPV+ cervical cancer cell growth *in vivo*.

**Figure 6.**
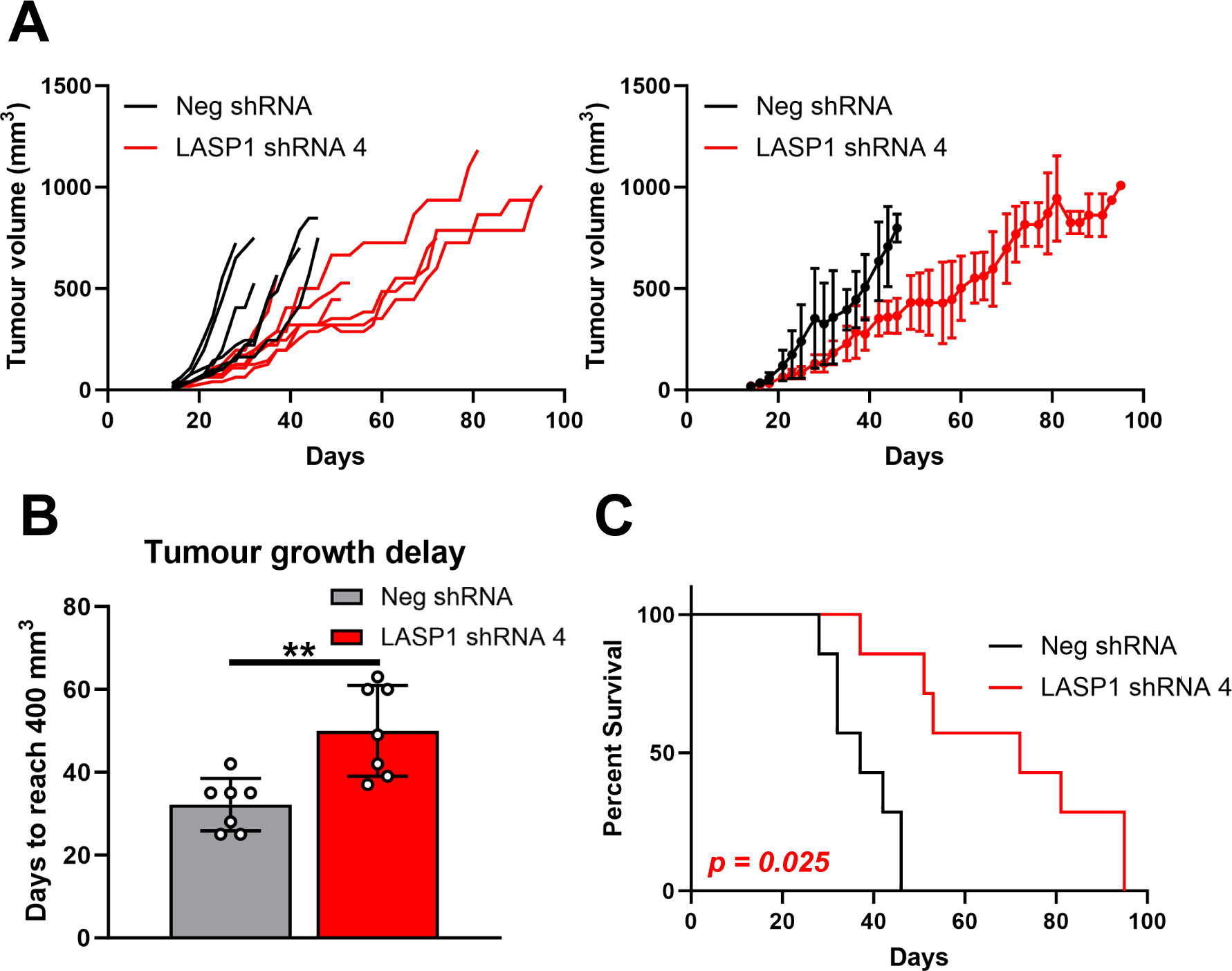
LASP1 depletion impairs HPV+ cervical cancer cell growth in an *in vivo* tumourigenicity model. **A)** Tumour growth curves for mice implanted with control HeLa cells (Neg shRNA) or HeLa LASP1 shRNA cells (LASP1 shRNA 1). Tumour volume was calculated using the formula V = 0.5*L*W2. Both individual curves for each replicate (left) and curves representing mean values +/− SD of seven mice per group (right) are displayed. **B)** Tumour growth delay, calculated as the period between injection of tumours and growth to a set volume (400 mm^3^). Bars represent means +/− SD of seven biological replicates with individual data points displayed. **C)** Survival curves of the mice in (**A**). Survival data were plotted using a Kaplan–Meier survival curve, and statistical significance was calculated using the log-rank test.

## Discussion

LASP1 was originally identified as a structural cytoskeletal protein with a scaffolding function (20,21,50). However, current data suggests that LASP1 is a potential oncogene in several cancers and has a diverse array of cellular functions, such as the regulation of cell signalling and gene expression (22,23,26,27,43,49). Additionally, LASP1 may have pathogenic roles beyond cancer, with a recent study showing a role for LASP1 in regulating adherens junction dynamics in inflammatory diseases such as arthritis (51). Here, we demonstrate that LASP1 functions as a proto-oncogene in HPV+ cervical cancer cells. Using a range of cervical cancer cell lines, precancerous lesions, and cervical cancer tissue microarrays, we showed that LASP1 is highly expressed in HPV+ cervical cancer cells. Depletion of LASP1, either transiently using siRNA or stably using shRNA, resulted in a significant proliferation defect in HPV+ cervical cancer cells, demonstrating the importance of LASP1 in driving the growth of these cancer cells. Interestingly, we did not observe a growth defect in HPV− cervical cancer cells upon LASP1 depletion; this suggests that LASP1-dependany may be HPV-specific. In a recent study, LASP1 depletion was shown to be detrimental to the proliferation of head and neck squamous cell carcinoma (HNSCC) cells, all of which were HPV− (43). It would therefore be interesting to see if the depletion of LASP1 in HPV+ HNSCC had a greater effect than on HPV− HNSCC cells. Of particular note, we observed a reduction in HPV E6 and E7 expression upon LASP1 depletion in HPV+ cervical cancer cells and in HPV18 containing keratinocytes. As expression of the viral oncoproteins are essential for the proliferation and survival of HPV+ cervical cancer cells, their depletion will have had an impact on the observed proliferation seen in our assays; however, it is important to note that over-expression of LASP1 in the absence of E6/E7 oncoproteins in HPV− C33A cells promoted proliferation. Therefore, we believe LASP1 drives proliferation independently of maintaining the expression of the HPV E6 and E7 and further experiments are needed to verify how LASP1 regulates their expression.

Through the use of primary keratinocytes containing the HPV16 or 18 genomes and expression constructs of the HPV oncogenes E6 and E7, our data shows that LASP1 expression was HPV E7 dependant. HPV E7 is a critical oncogene that activates a plethora of signalling pathways to drive cell proliferation and survival (14,15,19,52,53). Here, our data demonstrate that LASP1 constitutes a novel oncogenic target of HPV E7 in cervical cancer that may be an attractive target for cervical cancer and other HPV− associated cancers.

Previous studies have shown that HPV can manipulate miRNA networks to promote cellular transformation (16,38–40). miRNAs regulate many fundamental cellular functions, such as transcription, post-transcriptional modifications, and signal transduction (54). Data from our lab and others has shown that miR-203a, a well-studied tumour suppressor, is downregulated by HPV E6 and E7 (40,46–48). We have previously shown that the targeting of CREB1 is partially responsible for the proliferation defects observed upon miR-203 re-introduction in HPV+ cervical cancer cells (40). miR-203 can also regulate other targets in cervical cancer, including *VEGF* and *ZEB1* (55,56). Here, we demonstrate that miR-203 targets and represses LASP1 in HPV+ cervical cancer cells. miR-203 has been shown to regulate LASP1 in other squamous cell carcinomas, such as oesophageal cancer and HNSCC (43). Furthermore, a previous study demonstrated that the lncRNA CBR3-AS1 regulated LASP1 in cervical cancer via miR-3163; however, a functional assessment of the role LASP1 plays in cervical cancer was not examined further in this study (57). We demonstrate that miR-203 targets LASP1 in HPV+ cervical cancer cells and that the tumour suppressive effects of miR-203 introduction are ablated when LASP1 expression is restored. This suggests that LASP1 is a major miR-203 target in these cancers.

Our data further demonstrated that the SH3 domain of LASP1 is critical for its ability to drive proliferation in cervical cancer cells. LASP1 function is primarily mediated through multiple protein-protein interactions (21,49). Most of these interactions occur via either the LIM domain or the SH3 domain; however, the importance of either domain in mediating the pro-proliferative effects of LASP1 is not completely understood. Several important interactions between the LASP1 SH3 domain have been identified, including with LPP (50), Zo-2 (58) and Zyxin (59), and these are particularly important in regulating the scaffolding functions of LASP1 and its role in reorganisation of the actin cytoskeleton. Given our data, future studies aimed at understanding the importance of the LASP1 SH3 domain in the pro-proliferative effects observed in cervical cancer are warranted. Finally, we demonstrated that LASP1 expression is also critical for tumour growth *in vivo* by performing tumourigenicity assays. We observed significant delays to tumour growth in LASP1 KD cells which resulted a significant increase in survival, thus providing validation for our *in vitro* studies.

In conclusion, we present evidence that LASP1 plays an important role in cervical carcinogenesis. The viral oncoprotein E7 upregulates the expression of LASP1, which is a critical driver of proliferation in HPV+ cervical cancer cells. This is achieved through the repression on the microRNA miR-203, a potent tumour suppressor in squamous cell carcinomas. Importantly, we show that the SH3 domain of LASP1 is critical for the enhanced proliferation induced by LASP1 expression. A complete characterisation of the LASP1 interactome, particularly via the SH3 domain, is now warranted in order to determine if the functions of LASP1 can be targeted as a potential novel therapy in the treatment of HPV+ cervical cancers.

## Materials and Methods

### Cervical disease cytology samples

Cervical cytology samples were obtained from the Scottish HPV Archive (http://www.shine.mvm.ed.ac.uk/archive.shtml), a biobank of over 20,000 samples designed to facilitate HPV-associated research. The East of Scotland Research Ethics Service has given generic approval to the Scottish HPV Archive as a Research Tissue Bank (REC Ref 11/AL/0174) for HPV related research on anonymised archive samples. Samples are available for the present project though application to the Archive Steering Committee (HPV Archive Application Ref 0034). RNA was extracted from the samples using TRIzol® Reagent (ThermoFisher Scientific) and analysed as described.

### Plasmids, siRNA and miRNA products

Expression vectors for GFP-tagged HPV18 E6 and E7 have been described previously (17,18). pMSCV-N-HA-HPV18 E6-IRES-Puro, pMSCV-N-HA-HPV18 E7-IRES-Puro were previously described in (provided by Dr Elizabeth White, University of Pennsylvania, USA). The psiCheck2 plasmid was provided by Dr James Boyne (Leeds Becket University, UK). The *LASP1* 3’UTR was identified from NCBI data using the AceView program. It was subsequently amplified from HeLa cells and cloned into psiCheck2 using XhoI and NotI. For siRNA experiments, pools of four siRNAs specific to *LASP1* (FlexiTube GeneSolution GS3927) were purchased from Qiagen siRNA was used at a final concentration of 25 nM. For miRNA manipulations, hsa-miR-203a miRNA mimic (MIMAT0000264) and antagomir (MIMAT0031890) were obtained from ABM.

### Cell culture

HeLa (HPV18+ cervical epithelial adenocarcinoma cells), SW756 (HPV18+ cervical squamous carcinoma cells), SiHa (HPV16+ cervical squamous carcinoma cells), CaSki (HPV16+ cervical squamous carcinoma cells) and C33A (HPV− cervical squamous carcinoma) cells obtained from the ATCC were grown in DMEM supplemented with 10% FBS (ThermoFisher Scientific) and 50 U/mL penicillin/streptomycin (Lonza). HEK293TT cells were kindly provided by Prof Greg Towers (University College London (UCL)) and grown as above.

Neonate foreskin tissues were obtained from local General Practice surgeries and normal human keratinocytes (NHKs) were isolated under ethical approval no 06/Q1702/45. Cells were maintained in serum-free medium (SFM; Gibco) supplemented with 25 µg/mL bovine pituitary extract (Gibco) and 0.2 ng/mL recombinant EGF (Gibco). HPV18+ NHKs were generated as described previously [86]. N/Tert-1 and N/Tert-1+HPV16 cells were grown and maintained in K-SFM media containing 1% (vol/vol) penicillin-streptomycin mixture and 4 μg/ml hygromycin B ((36,62)).

NHKs and HPV18+ keratinocytes were grown in organotypic raft cultures by seeding onto collagen beds containing J2-3T3 fibroblasts (63). Once confluent, collagen beds were transferred onto metal grids in FCS-containing E media without EGF. The cells were allowed to stratify for 14 days before being fixed with 4% formaldehyde. Rafts were paraffin-embedded and 4 μm tissue sections prepared (Propath UK Ltd.). For analysis of LASP1, the formaldehyde-fixed raft sections were treated with the sodium citrate method of antigen retrieval. Briefly, sections were boiled in 10 mM sodium citrate with 0.05% Tween-20 for 10 minutes. Sections were incubated with a polyclonal antibody against LASP1 (G-7; sc-374059, Santa Cruz Biotechnology (SCBT)) and immune complexes visualised using Alexa 594 secondary antibodies (Invitrogen). Nuclei were counterstained with DAPI and mounted in Prolong Gold (Invitrogen).

All cells were cultured at 37 °C and 5% CO2 were negative for mycoplasma during this investigation. Where appropriate, cell identity was confirmed by STR profiling.

### Mammalian cell transfection

Transient transfections were performed using Lipofectamine 2000 (ThermoFisher Scientific) at a ratio of nucleic acid:Lipofectamine 2000 of 1:2 for both DNA and siRNA. Transfections were performed overnight in OptiMEM I Reduced Serum Media (ThermoFisher Scientific). Subsequent analyses were performed at 48 hours (DNA) or 72 hours (siRNA) post-transfection.

### Generation of stable cell lines

HeLa and SiHa LASP1 Knock down (KD) cell lines were generated using LASP-1 shRNA (h) Lentiviral Particles (sc-105607-V, SCBT). To perform lentiviral transduction, culture media was removed from cells seeded 24 hours earlier and replaced with virus-containing media. Cells were incubated overnight before removing virus and replacing with complete DMEM. At 48 hours post-transduction, cells were passaged as appropriate and treated with 1 μg/mL puromycin in culture media for 48 hours to select for transduced cells. To generate monoclonal KD cell lines, polyclonal stocks were diluted to 1 cell per well manually in a 96 well plate and surviving cells were screened for sufficient knockdown of target gene via RT-qPCR. Two separate clones were generated for each cell line. HPV− C33A cells stably expressing HPV18 E6 or E7 were generated as previously described (16).

### Western blot analysis

Equal amounts of protein from cell lysates were seperated using 8-15% SDS-polyacrylamide gels as appropriate. Separated proteins were transferred to Hybond™ nitrocellulose membranes (GE Healthcare) using a semi-dry method (Bio-Rad Trans-Blot® Turbo™ Transfer System). Membranes were blocked in 5% skimmed milk powder in tris-buffered saline-0.1% Tween 20 (TBS-T) for 1 hour at room temperature before probing with antibodies specific for HPV16 E6 (GTX132686, GeneTex, Inc.), HPV16 E7 (ED17: sc-6981, SCBT), HPV18 E6 (G-7: sc-365089, SCBT), HPV18 E7 (8E2: ab100953, abcam), HA (3724, Cell Signalling Technology (CST)), LASP1 (G-7; sc-374059; SCBT) and GAPDH (G-9: sc-365062, SCBT). Primary antibody incubations were performed overnight at 4 °C. The appropriate HRP-conjugated secondary antibodies (Jackson ImmunoResearch) were used at a 1:5000 dilution. Blots were visualised using ECL reagents and CL-XPosure™ film (ThermoFisher Scientific). A minimum of three biological repeats were performed in all cases and representative blot images are displayed.

### RNA extraction and reverse transcription-quantitative PCR (RT-qPCR)

Total RNA was extracted from cells using the E.Z.N.A.® Total RNA Kit I (Omega Bio-Tek) following the provided protocol for RNA extraction from cultured cells. The concentration of eluted RNA was determined using a NanoDrop™ One spectrophotometer (ThermoFisher Scientific). RT-qPCR was performed using the GoTaq® 1-Step RT-qPCR System (Promega) with an input of 50 ng RNA. Reactions were performed using a CFX Connect Real-Time PCR Detection System (BioRad) with the following cycling conditions: reverse transcription for 10 min at 50 °C; reverse transcriptase inactivation/polymerase activation for 5 min at 95 °C followed by 40 cycles of denaturation (95 °C for 10 sec) and combined annealing and extension (60 °C for 30 sec). Data was analysed using the ΔΔCt method (64).

### Tissue microarray and immunohistochemistry

A cervical cancer tissue microarray (TMA) containing 39 cases of cervical cancer and 9 cases of normal cervical tissue (in duplicate) was purchased from GeneTex, Inc. (GTX21468). Slides were deparaffinised in xylene, rehydrated in a graded series of ethanol solutions and subjected to antigen retrieval in citric acid. Slides were blocked in normal serum and incubated in primary antibody against LASP1 (G-7; sc-374059; SCBT) overnight at 4 °C. Slides were then processed using the VECTASTAIN^®^ Universal Quick HRP Kit (PK-7800; Vector Laboratories) as per the manufacturer’s instructions. Immunostaining was visualised using 3,3’-diaminobenzidine (Vector^®^ DAB (SK-4100; Vector Laboratories)). Images were taken using an EVOS® FL Auto Imaging System (ThermoFisher Scientific) at 20x magnification. Protein quantification was automated using ImageJ with the IHC Profiler plug-in (65,66). Histology scores (H-score) were calculated based on the percentage of positively stained tumour cells and the staining intensity grade. The staining intensities were classified into the following four categories: 0, no staining; 1, low positive staining; 2, positive staining; 3, strong positive staining. H-score was calculated by the following formula: (3 x percentage of strong positive tissue) + (2 x percentage of positive tissue) + (percentage of low positive tissue), giving a range of 0 to 300.

### Luciferase reporter assays

Cells were transfected with plasmids expressing the appropriate firefly luciferase reporter. A *Renilla* luciferase reporter construct (pRLTK) was used as an internal control for transfection efficiency. Samples were lysed in passive lysis buffer (Promega) and activity measured using a dual-luciferase reporter assay system (Promega). All assays were performed in triplicate, and each experiment was repeated a minimum of three times.

### Proliferation assays

For cell growth assays, cells were trypsinised after treatment as necessary and reseeded. Cells were subsequently harvested every 24 hours and manually counted using a haemocytometer.

For colony formation assays, cells were trypsinised after treatment as required and reseeded at 500 cells/well. Once visible colonies were noted, cells were fixed and stained in crystal violet staining solution (1% crystal violet, 25% methanol) for 15 min at room temperature. Plates were washed thoroughly with water to remove excess crystal violet and colonies counted manually.

For soft agar assays, 60 mm cell culture plates were coated with a layer of complete DMEM containing 0.5% agarose. Simultaneously, cells were trypsinised after treatment as required and resuspended at 1000 cells/mL in complete DMEM containing 0.35% agarose and added to the bottom layer of agarose. Once set, plates were covered with culture media and incubated for 14-21 days until visible colonies were observed. Colonies were counted manually.

### *In vivo* tumourigenicity study

Female 6-8 week old SCID mice were purchased from Charles River Laboratories. All animal work was carried out under project license PP1816772. HeLa cells stably expressing either a non-targeting shRNA (Neg shRNA) or a LASP1-specific shRNA were harvested, pelleted, and resuspended in sterile PBS. Seven mice were used per experimental group, with each injected subcutaneously with 5 x 10^5^ cells in 50 µL PBS. Once palpable tumours had formed (∼10 days), measurements for both groups were taken thrice weekly. After tumours reached 10 mm in either dimension, mice were monitored daily. Mice were sacrificed once tumours reached 15 mm in any dimension. No toxicity, including significant weight loss, was seen in any of the mice. Tumour volume was calculated with the formula V = 0.5*L*W^2^.

### Database analysis

Patient overall survival (OS) data were downloaded from the Supplemental Information of a pan-cancer clinical study (67). OS curves were obtained using Kaplan-Meier method with expression cut off determined using KMPlotter (68) and were compared using the log-rank test. The Cox proportional hazards model was used to estimate Hazard Ratios (HRs) with 95% Confidence Intervals (CIs).

Microarray data was obtained from GEO database accession number GSE6791 (32), GSE7803 (33), GSE9750 (34) and GSE63514 (35).

### Statistical analysis

All *in vitro* experiments were performed a minimum of three times, unless stated otherwise. The sample size for the *in vivo* study was selected by assuming an expected mean survival of the control animal group of approximately 40 ± 5 days (based on previous experiments in the subcutaneous SCID/HeLa flank model (19)). Data was analysed using a two-tailed, unpaired Student’s t-test performed using GraphPad PRISM 9.2.0 software, unless stated otherwise. Kaplan-Meier survival data was analysed using the log-rank (Mantel-Cox) test.

## Acknowledgements

We thank the Scottish HPV Investigators Network (SHINE), Prof Sheila Graham (University of Glasgow), Dr David Millan (University of Glasgow) and Prof Nick Coleman (University of Cambridge) for providing HPV positive patient samples.

## Funding information

Work in the Macdonald lab is supported by Medical Research Council (MRC) funding (MR/ K012665, MR/S001697/1 and MR/X009564/1). ELM was supported by the Wellcome Trust (1052221/Z/14/Z and 204825/Z/16/Z) and the University of Sussex. MRP was funded by a Biotechnology and Biological Sciences Research Council (BBSRC) studentship (BB/M011151/1). ASM was funded by the Erasmus+ programme. ELR, ALT and JED were supported by the Wellcome Trust (105210/Z/14/Z and 102174/B/13/Z). JAS was funded by a Faculty of Biological Sciences, University of Leeds Scholarship. J.E.L. was funded by a CRUK Program Grant (C57233/A22356). MW was supported by a Leeds-China Scholarship. CDJ and IMM are supported by NIH grant R01DE029471. AS is funded by CRUK (C50189/A29039). The funders had no role in study design, data collection and analysis, decision to publish, or preparation of the manuscript.

### Author Contributions

Conceptualisation (ELM, AM); Formal analysis (MRP, ASM, JAS, DE, ELM); Investigation (MRP, ASM, ELR, JAS, DE, ALT, CWW, JED, MW, ELM); Resources (JED, CDJ, JEL, IMM); Funding acquisition (JEL, IMM, AS, ELM, AM); Project administration (AS, ELM, AM); Supervision (JEL, IMM, AS, ELM, AM); Writing – original draft (ELM); Writing – review & editing (all authors)

---

**Supp Figure 1.**
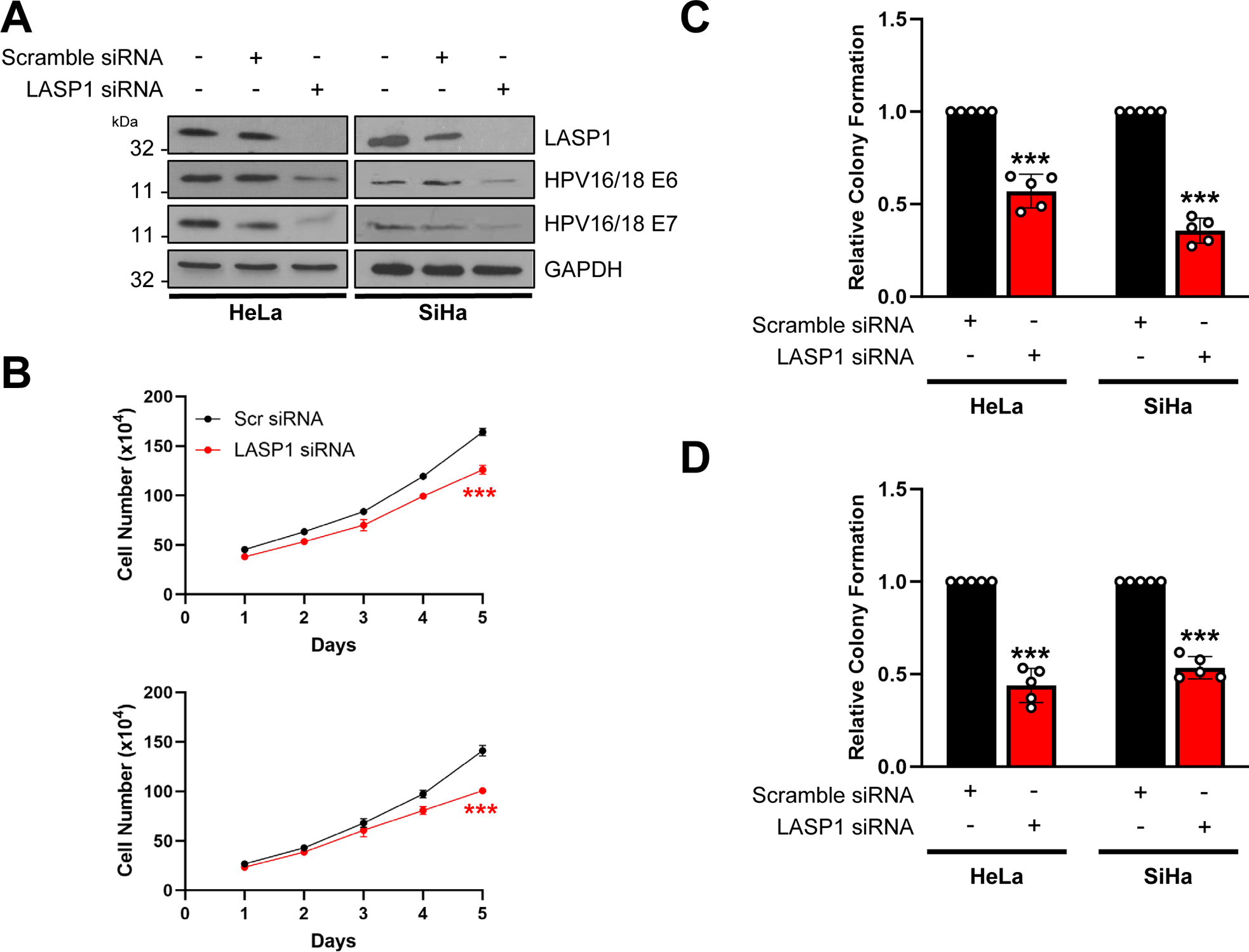
Transient LASP1 depletion inhibits proliferation in HPV+ cervical cancer cells. **A)** Representative western blot of LASP1 protein expression in HeLa and SiHa cells after LASP1 depletion with a pool of four specific siRNAs. Lysates were probed for LASP1, HPV E6 and HPV E7. GAPDH was used as a loading control. **B)** Growth curve assay in HeLa and SiHa cells after LASP1 depletion with a pool of four specific siRNAs. Colony formation assay in HeLa and SiHa cells after LASP1 depletion with a pool of four specific siRNAs. **D)** Soft Agar assay in HeLa and SiHa cells after LASP1 depletion with a pool of four specific siRNAs. Error bars represent the mean +/− standard deviation of a minimum of three biological repeats unless otherwise stated. *ns –* not significant, \**p* < 0.05, \*\**p* < 0.01, \*\*\**p* < 0.001 (Student’s *t*-test).

**Supp Figure 2.**
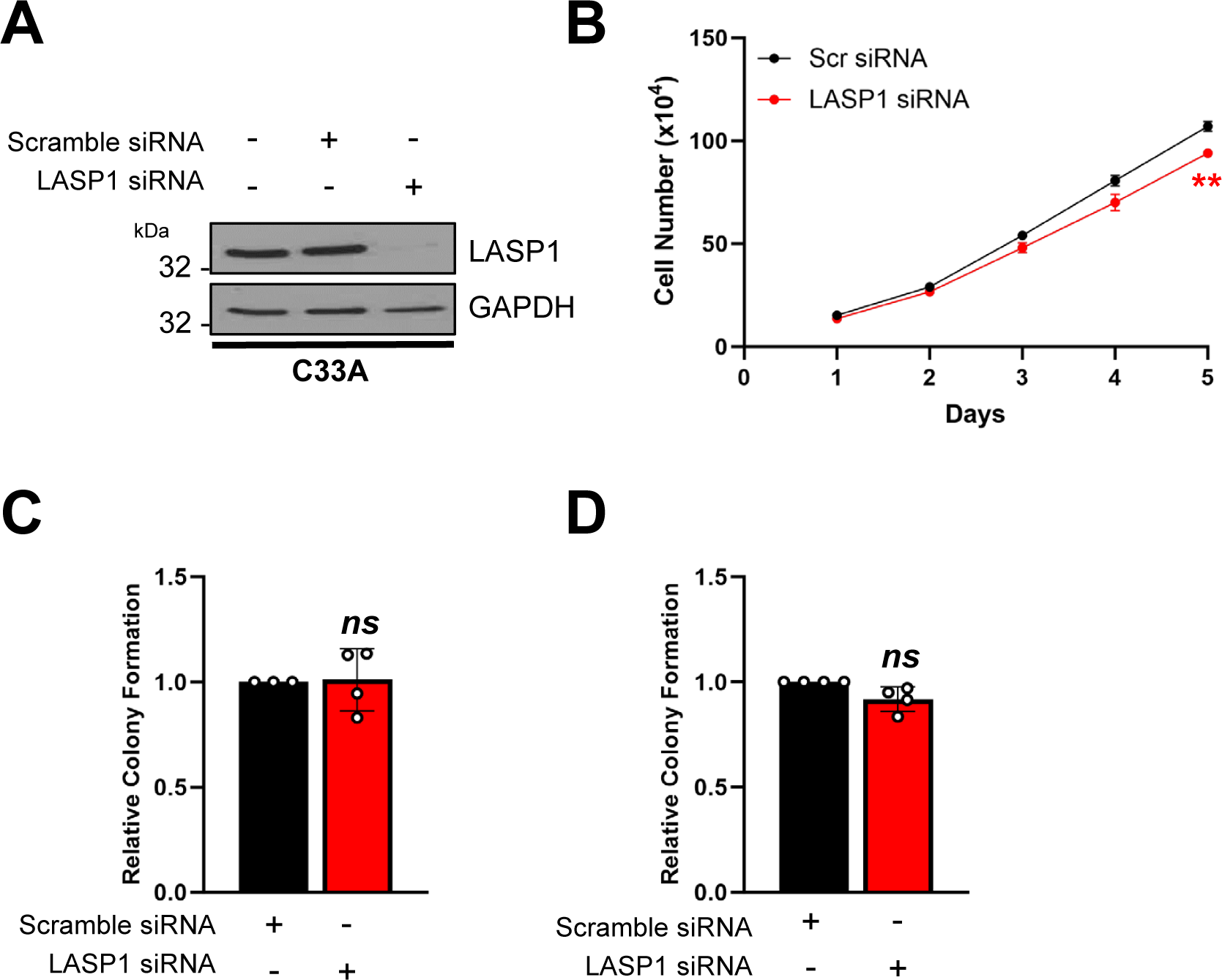
Transient LASP1 depletion has minimal impact on the proliferation of HPV− cervical cancer cells. **A)** Representative western blot of LASP1 protein expression in C33A cells after LASP1 depletion with a pool of four specific siRNAs. Lysates were probed for LASP1 and GAPDH was used as a loading control. **B)** Growth curve assay in C33A cells after LASP1 depletion with a pool of four specific siRNAs. **C)** Colony formation assay in C33A cells after LASP1 depletion with a pool of four specific siRNAs. **D)** Soft Agar assay in C33A cells after LASP1 depletion with a pool of four specific siRNAs. Error bars represent the mean +/− standard deviation of a minimum of three biological repeats unless otherwise stated. *ns –* not significant, \**p* < 0.05, \*\**p* < 0.01, \*\*\**p* < 0.001 (Student’s *t*-test).

**Supp Figure 3.**
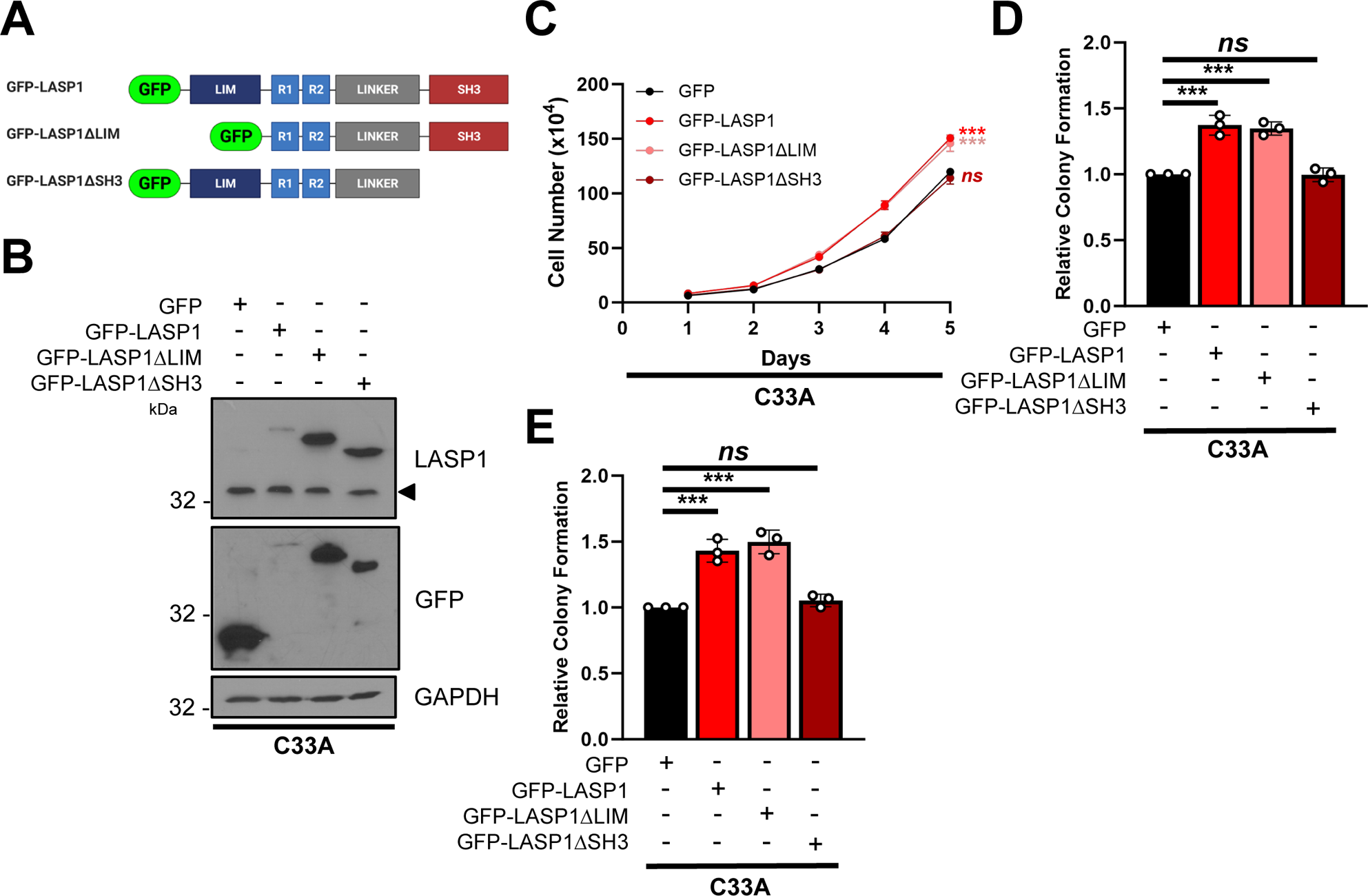
LASP1 promotes proliferation in HPV− cervical cancer cells in an SH3-domain dependent manner. **A)** Schematic showing the LASP1 mutant constructs used in the study. **B)** Representative western blot of LASP1 mutants in C33A cells. GFP-LASP1, GFP-LASP1ΔLIM and GFP-LASP1ΔSH3 expression were confirmed using GFP and LASP1 antibodies. Arrow indicates endogenous LASP1 expression. GAPDH was used as a loading control. **C)** Growth curve assay in C33A cells expressing GFP-LASP1, GFP-LASP1ΔLIM and GFP-LASP1ΔSH3. **D)** Colony formation assay in C33A cells expressing GFP-LASP1, GFP-LASP1ΔLIM and GFP-LASP1ΔSH3. **E)** Soft Agar in C33A cells expressing GFP-LASP1, GFP-LASP1ΔLIM and GFP-LASP1ΔSH3. Error bars represent the mean +/− standard deviation of a minimum of three biological repeats unless otherwise stated. *ns –* not significant, \**p* < 0.05, \*\**p* < 0.01, \*\*\**p* < 0.001 (Student’s *t*-test).

